# Identification of a novel putative interaction partner of the nucleoporin ALADIN

**DOI:** 10.1101/064766

**Authors:** Ramona Jühlen, Dana Landgraf, Angela Huebner, Katrin Koehler

**Affiliations:** Klinik und Poliklinik für Kinder- und Jugendmedizin, Medizinische Fakultät Carl Gustav Carus, Technische Universität Dresden, Germany

**Keywords:** ALADIN, Cell division, Nuclear pore complex, PGRMC2, Triple A syndrome

## Abstract

It has been shown that the nucleoporin ALADIN employs a significant role in the redox homeostasis of the cell but the function in steroidogenesis contributing to adrenal atrophy in triple A syndrome remains largely unknown. In an attempt to identify new interaction partners of ALADIN, co-immunoprecipitation followed by proteome analysis was conducted in different expression models using the human adrenocortical tumour cell line NCI-H295R. Our results suggest an interaction of ALADIN with the microsomal protein PGRMC2. PGRMC2 is shown to be activity regulator of CYP P450 enzymes and therefore, to be a possible target for adrenal dysregulation in triple A syndrome. We show that there is a sexual dimorphism regarding the expression of *Pgrmc2* in adrenals and gonads of WT and *Aaas* KO mice. Female *Aaas* KO mice are sterile due to delayed oocyte maturation and meiotic spindle assembly. A participation in meiotic spindle assembly confirms the recently investigated involvement of ALADIN in mitosis and emphasises an interaction with PGRMC2 which is a regulator of cell cycle. By identification of a novel interaction partner of ALADIN we provide novel aspects for future research of the function of ALADIN during cell cycle and for new insights into the pathogenesis of triple A syndrome.

## Summary statement

In our study we report the interaction of the microsomal integral membrane protein progesterone receptor membrane compartment 2 with the nucleoporin ALADIN.

## Introduction

The triple A syndrome (MIM#231550) is an autosomal recessive disease manifesting with the triad of ACTH-resistant adrenal insufficiency, achalasia of the stomach cardia and alacrima in combination with progressive neurological impairment (Allgrove et al., 1978). The syndrome is caused by mutations in the *AAAS* (achalasia-adrenal insufficiency-alacrima syndrome) gene encoding the protein ALADIN (**al**acrima-**a**chalasia-a**d**renal **in**sufficiency neurologic disorder) (Handschug et al., 2001; Tullio-Pelet et al., 2000). ALADIN is ubiquitously expressed, but shows enhanced levels in neuroendocrine and gastrointestinal structures; tissues which are most affected in triple A patients (Handschug et al., 2001).

ALADIN is a scaffold nucleoporin (NUP) anchored within the nuclear pore complex (NPC) by the transmembrane NUP NDC1 (nuclear division cycle 1 homologue (*S. cerevisiae*)) (Kind et al., 2009; Yamazumi et al., 2009). It belongs to the group of barely exchangeable NUPs and therefore seems to be involved in building the structural scaffold backbone of the complex at the nuclear membrane (Rabut et al., 2004). Over the last years it has been shown that NUPs have fundamental functions in cell biology, especially beyond nucleo-cytoplasmic transport (Fahrenkrog, 2014; Nofrini et al., 2016).

Our group has reported that ALADIN is involved in the oxidative stress response of fibroblasts and adrenocortical cells but the role of ALADIN in adrenal steroidogenesis contributing to the adrenal phenotype in triple A patients is largely unknown (Jühlen et al., 2015; Kind et al., 2010; Koehler et al., 2013; Storr et al., 2009). Recently, we showe/d that a depletion of ALADIN in adrenocortical carcinoma cells leads to an alteration in glucocorticoid and androgenic steroidogenesis and a diminished redox homeostasis (Jühlen et al., 2015). Our results described in this article propose an interaction of ALADIN with the microsomal integral membrane protein **p**ro**g**esterone **r**eceptor **m**embrane **c**ompartment **2** (PGRMC2). PGRMC2 belongs to the group of membrane-associated progesterone receptors (MAPRs). These receptors are restricted to the ER and are thought to act on mitosis while localising to the somatic spindle apparatus and to regulate the activity of some CYP P450 enzymes (e.g. CYP21A2) (Keator et al., 2012; Peluso et al., 2014; Wendler and Wehling, 2013). By the attempt to identify new interaction partners of ALADIN we aimed to clarify the cellular functions of ALADIN at the NPC and to explain the mechanisms which contribute to the adrenal insufficiency in triple A syndrome. Our observations give the basis for further research on the association between ALADIN and PGRMC2 and about the function of ALADIN during cell cycle and steroidogenesis beyond nucleo-cytoplasmic transport.

## Results

### PGRMC2 precipitates with ALADIN in an exogenous and endogenous ALADIN adrenal cell expression model

Co-IP was conducted in NCI-H295R cells either expressing endogenous ALADIN or additionally exogenous GFP-ALADIN.

We performed mass spectrometry analyses of bound fractions of GFP(-ALADIN) and ALADIN co-IP. In the exogenous GFP-ALADIN expression model sufficient ALADIN peptides could be identified (**Fig. 1A**), the analysis of endogenous ALADIN co-IP resulted in less detected peptides (**Fig. 1B**). Despite distinct methodological optimisation procedures using different protocols and antibodies, endogenous ALADIN co-IP had a low yield and measurement of bound fractions using mass spectrometry was more difficult to process. All proteins identified in mass spectrometry in GFP-ALADIN co-IP and ALADIN co-IP but which were not found in the specific control pulldown assays are presented in the supplementary data (**Tables S1 and S2**).

**Figure 1.**
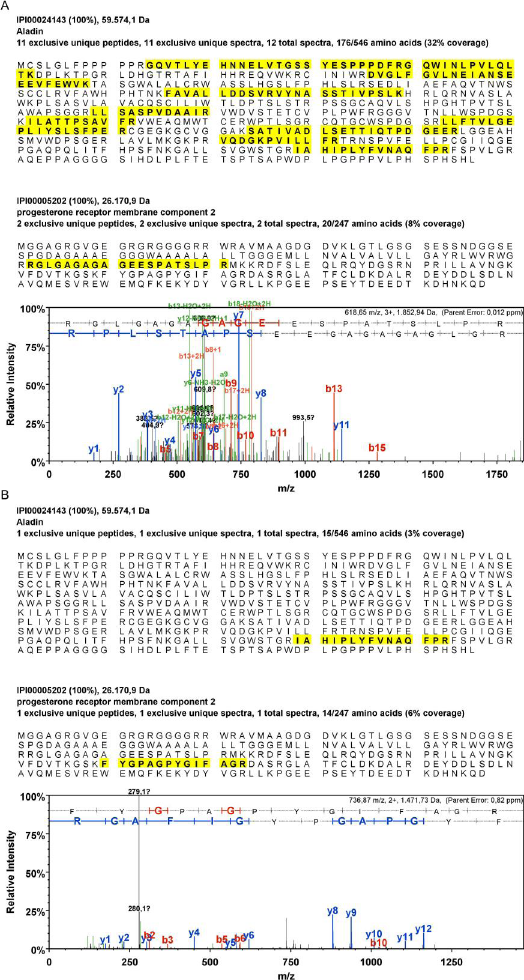
PGRMC2 was identified in mass spectrometry after (A) GFP-(ALADIN) and (B) ALADIN pulldown. Identified exclusive unique peptides (yellow) (number of different amino acid sequences, regardless of any modification, that are associated only with this protein) are shown of ALADIN and progesterone receptor membrane compartment 2 (PGRMC2). For PGRMC2 also annotated spectra are shown: (A) after GFP(-ALADIN) co-IP of whole cell lysates of GFP-ALADIN expressing NCI-H295R cells and (B) after ALADIN co-IP of whole cell lysates of NCI-H295R cells.

PGRMC2 was simultaneously identified by mass spectrometry analysis in co-IP of GFP-ALADIN using GFP-Trap_A agarose beads and in co-IP of ALADIN using anti-ALADIN coupled to Protein G UltraLink resin sepharose beads. Exclusive unique peptides of ALADIN and PGRMC2 detected in mass spectrometry after GFP-ALADIN and ALADIN pulldown are shown in **Fig. 1A** and **1B**, respectively.

We additionally confirmed the identification of PGRMC2 in GFP-ALADIN and ALADIN co-IP by Western Blot. Successful GFP-ALADIN pulldown in lysates of NCI-H295R cells stably expressing GFP-ALADIN (86 kDa) is presented in **Fig. 2A**. In accordance with our mass spectrometry results PGRMC2 (24 kDa) could also be detected after GFP-ALADIN pulldown (**Fig. 2A**, arrow). The negative control remained empty.

**Figure 2.**
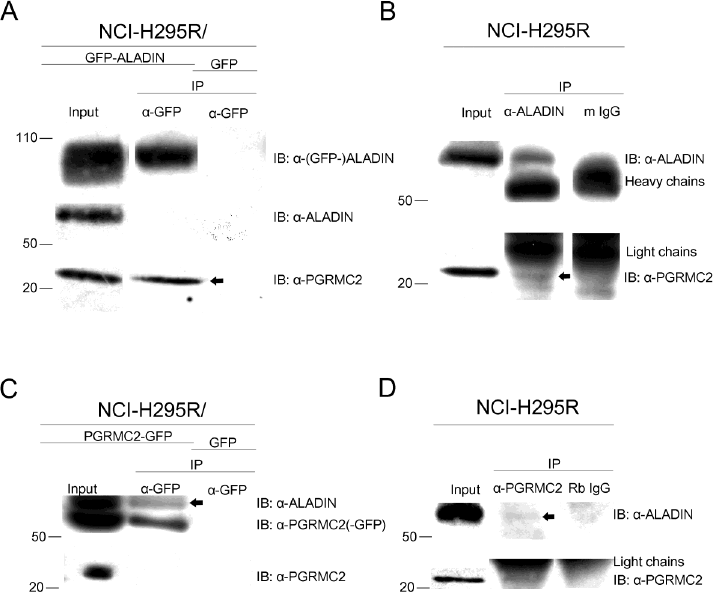
PGRMC2 interacts with ALADIN determined by IP-Western and reciprocal IP-Western assays. (A) Whole cell lysates of GFP-ALADIN and GFP expressing NCI-H295R cells were used and GFP pulldown performed followed by Western Blot with indicated antibodies. PGRMC2 (24 kDa) (arrow) could be detected after GFP-ALADIN (86 kDa) pulldown. (B) Whole cell lysates of NCI-H295R cells were used for ALADIN pulldown and normal mouse IgGs (m IgG) as negative control followed by Western Blot with indicated antibodies. PGRMC2 (arrow) precipitated in endogenous ALADIN (59 kDa) pulldown. (C) Whole cell lysates of PGRMC2-GFP and GFP expressing NCI-H295R cells were used and reciprocal GFP pulldown was performed followed by Western Blot with indicated antibodies. ALADIN (arrow) could be detected after PGRMC2-GFP (51 kDa) pulldown. (D) Whole cell lysates of NCI-H295R cells were used for PGRMC2 pulldown and normal rabbit IgGs (Rb IgG) as negative control followed by Western Blot with indicated antibodies. ALADIN (arrow) slightly but visibly precipitated after PGRMC2 pulldown.

Successful endogenous ALADIN pulldown is shown in **Fig. 2B**. ALADIN (59 kDa) was found in the bound fraction of the ALADIN co-IP but the negative control using normal mouse IgG was shown to be empty. In accordance to our result after GFP-ALADIN pulldown, PGRMC2 precipitated in endogenous ALADIN pulldown with no unspecific interaction in the negative control (**Fig. 2B**, arrow).

### ALADIN precipitates with PGRMC2 in an exogenous and endogenous PGRMC2 adrenal cell expression model

In order to further evaluate a possible interaction between the nucleoporin ALADIN and microsomal PGRMC2 we conducted reciprocal co-IP assays.

Reciprocal pulldowns were done in NCI-H295R cells transiently expressing exogenous PGRMC2-GFP and endogenous PGRMC2.

Efficient pulldown of (PGRMC2-)GFP in lysates of NCI-H295R cells transiently expressing PGRMC2-GFP (51 kDa) is presented in **Fig. 2C**. Confirming our previous results identifying a possible interaction between ALADIN and PGRMC2, ALADIN could be detected after PGRMC2-GFP pulldown (**Fig. 2C**, arrow). The negative control was empty.

PGRMC2 pulldown is shown in **Fig. 2D**. PGRMC2 was successfully detected in the bound fraction of the PGRMC2 co-IP and the negative control using rabbit IgG remained empty. According to our results after PGRMC2-GFP pulldown, ALADIN slightly but visibly precipitated after PGRMC2 pulldown with no unspecific interaction in the negative control (**Fig. 2D**, arrow).

### Localisation of ALADIN and PGRMC2 in adrenal cells using different expression models

Further evidence of possible co-localisation of ALADIN and PGRMC2 is given after immunofluorescent staining. Cells expressing GFP-ALADIN and PGRMC2-GFP were used to verify unspecific staining of anti-ALADIN and anti-PGRMC2 in NCI-H295R cells. Staining was done using anti-ALADIN, anti-PGRMC2 and anti-NPC proteins (mAb414).

Immunostaining with mAb414 in all adrenal cell expression models gave a thin circle around the nucleus indicating punctuate localisations of NPCs (**Fig. 3**).

**Figure 3.**
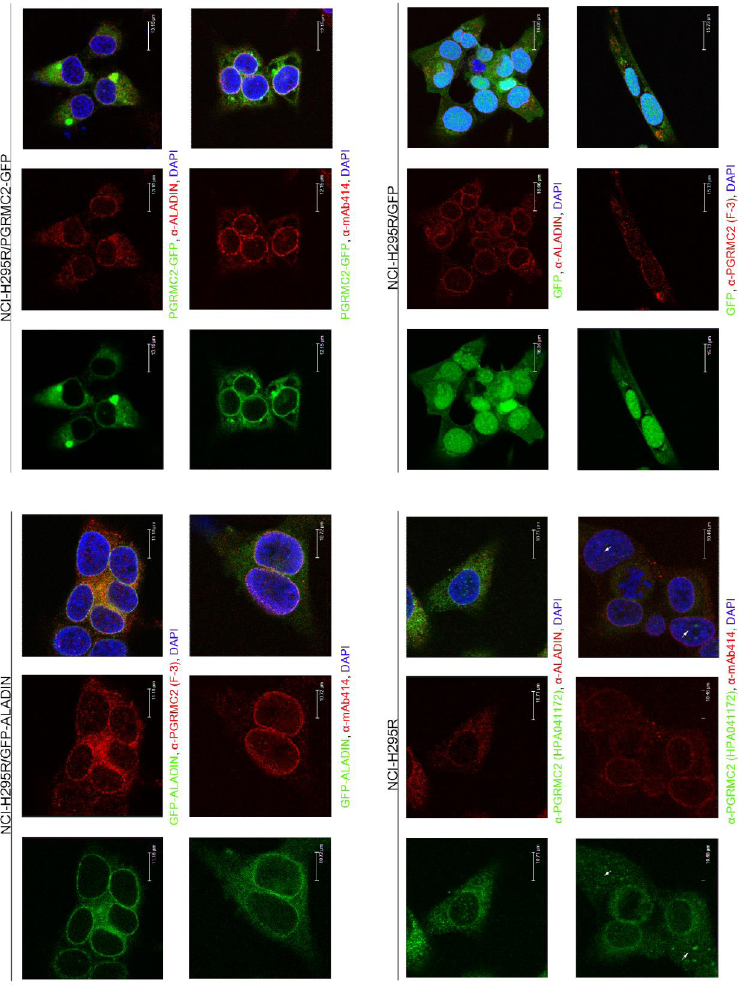
ALADIN and PGRMC2 localise to the perinuclear space in human adrenocortical carcinoma cells. NCI-H295R/GFP-ALADIN, NCI-H295R/PGRMC2-GFP, NCI-H295R and NCI-H295R/GFP cells were stained with anti-ALADIN, anti-PGRMC2, anti-NPC proteins (mAb414) and DAPI.

Immunofluorescent staining of the nucleoporin ALADIN appeared at the nuclear envelope at the proximity of NPCs in all adrenal cell expression models. In the exogenous GFP-ALADIN cell model the fusion protein was correctly targeted to the nuclear envelope and did not accumulate to a greater extent in the cytoplasm. We showed that ALADIN almost completely co-localises with anti-NPC proteins (mAb414) immunostaining at the nuclear envelope; substantially verifying the localisation of ALADIN at the nuclear pore (**Fig. 3**).

In immunofluorescent staining the microsomal protein PGRMC2 localised to the central ER but also revealed a patchy and punctuate staining pattern around the nucleus to the perinuclear space between nuclear envelope and ER in all adrenal cell expression models (**Fig. 3**). The same PGRMC2 immunostaining pattern was observed in human cervical carcinoma (HeLa) (**Fig. S1A**) and human fibroblasts (**Fig. S1B**). In the exogenous adrenal cell model the PGRMC2-GFP fusion protein was still correctly targeted to the central ER and perinuclear space. In the staining with the anti-PGRMC2 in NCI-H295R cells we could also observe nuclear staining in some adrenal cells (**Fig. 3**, arrow). Nuclear localisation of PGRMC2 was absent in the PGRMC2-GFP adrenal cell expression model. Co-localisation between mAb414 and PGRMC2 or ALADIN and PGRMC2 was not complete but showed positivity in the perinuclear space and in the nuclear membrane in all cell expression models (**Fig. 3**).

### The expression of PGRMC2 is not affected after AAAS knock-down in human adrenal cells

To test if *PGRMC2* expression is affected when ALADIN is down-regulated we used the inducible NCI-H295R1-TR cells with *AAAS* knock-down shRNA or scrambled shRNA as negative control (Jühlen et al., 2015).

We could not find an alteration on PGRMC2 mRNA level after induction of ALADIN depletion by doxycycline in NCI-H295R1-TR cells in at least ten triplicate experiments (**Fig. S2**).

### Pgrmc2 exhibits a sexual dimorphism in adrenals and gonads of WT and Aaas KO mice

In order to examine the expression of *Pgrmc2* in WT and *Aaas* KO mice we looked at the adrenals and gonads using Taq Man analysis and Western Blot (**Fig. 4A**).

**Figure 4.**
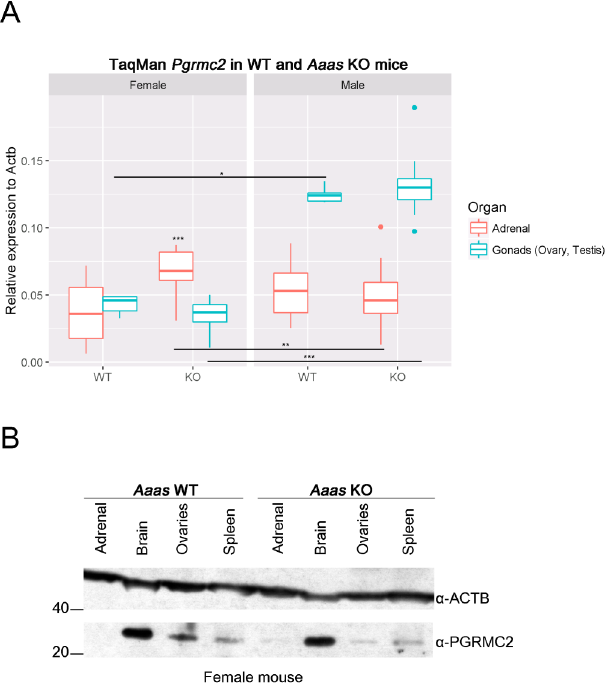
*Pgrmc2* has a sexual dimorphic role in mice and ALADIN KO in female mice leads to an alteration in PGRMC2. (A) Total RNA was isolated from dissected adrenals and gonads of WT and *Aaas* KO mice. P-values: * P < 0.05, ** P < 0.01, *** P < 0.001. Significant differences were measured with unpaired Wilcoxon-Mann-Whitney U-test. Boxplot widths are proportional to the square root of the samples sizes. Whiskers indicate the range outside 1.5 times the inter-quartile range (IQR) above the upper quartile and below the lower quartile. Outliers were plotted as dots. (B) Total protein was isolated from dissected adrenals, brain, ovaries and spleen of female WT and *Aaas* KO mice followed by Western Blot with indicated antibodies.

Indeed, we found that *Pgrmc2* exploited a sexual dimorphism in female and male WT and *Aaas* KO mice: the expression in testes was significantly higher independent of genotype compared to female ovaries, in female KO adrenals the expression was significantly higher compared to male KO adrenals (**Fig. 4A**).

Interestingly, in female adrenals the depletion of ALADIN leads to a significant increase in *Pgrmc2* expression whereas in female ovaries a decrease compared to WT ovaries was observed (**Fig. 4A**). The expression of *Pgrmc2* was not altered in male *Aaas* KO adrenals or testes compared to male WT organs in at least four triplicate experiments (**Fig. 4A**).

To examine our findings in female WT and *Aaas* KO mice on PGRMC2 RNA level, we conducted Western Blot of several female murine WT and *Aaas* KO tissues, i.e. adrenals, brain, ovaries and spleen (**Fig. 4B**). We could confirm our results on PGRMC2 RNA level and could show an increase in PGRMC2 protein in female adrenals of *Aaas* KO mice compared to female adrenals of WT mice (**Fig. 4B**). Furthermore, in ovaries of *Aaas* KO mice PGRMC2 protein was diminished compared to ovaries of WT mice.

## Discussion

The exact role of the nucleoporin ALADIN at the NPC and its involvement in steroidogenesis leading to the characteristic adrenal atrophy in triple A syndrome remains largely unknown. We and others have provided proofs of the involvement of ALADIN in the oxidative stress response of the cell (Jühlen et al., 2015; Kind et al., 2010; Koehler et al., 2013; Prasad et al., 2013; Storr et al., 2009). Recently, we could show that a depletion of ALADIN in adrenocortical carcinoma cells leads to an alteration in glucocorticoid and androgenic steroidogenesis (Jühlen et al., 2015).

Despite the reported interaction between ALADIN and ferritin heavy chain 1 no other interaction partner which would lead to the identification of a plausible function and signal transduction of ALADIN in the cell is known so far (Storr et al., 2009).

In an attempt to identify new interaction partners of ALADIN co-IP analyses showed that PGRMC2 precipitated with ALADIN. To verify the identified association between ALADIN and PGRMC2 reciprocal IP was conducted. Our results showed a co-precipitation of ALADIN with PGRMC2. By different co-IP approaches using exogenous and endogenous expression systems in human adrenal cells we can show for the first time that the nucleoporin ALADIN associates in a complex with the microsomal protein PGRMC2.

PGRMC2 belongs to the MAPR family. MAPRs are restricted to the ER and are thought to regulate the activity of CYP P450 enzymes. The first identified MAPR, PGRMC1, gained widespread attention (Falkenstein et al., 1996). PGRMC1 is a cytochrome-related protein with several implications in cancer (Clark et al., 2016; Falkenstein et al., 1996; Kabe et al., 2016)

In this work, PGRMC2 was found to interact with the nucleoporin ALADIN. PGRMC2 is barely investigated compared to its homologue PGRMC1. It is known that PGRMC2 alters activity of CYP3A4 as possible electron donor, and binds CYP21A2, most likely through its cytochrome b5-similar heme-binding domain (Albrecht et al., 2012; Wendler and Wehling, 2013).

Visualising PGRMC2 in the cell using immunofluorescence and confocal microscopy PGRMC2 appeared at the central ER and interestingly, at the nuclear envelope and the perinuclear ER. We detected that PGRMC2 co-localises with ALADIN and with different FG-repeat NUPs (stained with anti-NPC proteins (mAb414)) to the nuclear envelope and the perinuclear ER. Taken together, our results in immunofluorescence microscopy using different ALADIN and PGRMC2 adrenal cell expression systems provide a basis for future research of how ALADIN and PGRMC2 possibly associate in a complex close to the nuclear envelope and what the molecular function of this association would be.

Furthermore, we present that ALADIN depletion did not result in diminished PGRMC2 expression on mRNA level *in vitro* in adrenocortical carcinoma cells. However, we found that *Pgrmc2* has a sexual dimorphic role in adrenals and gonads of WT and *Aaas* KO mice and of note, ALADIN depletion leads to an alteration in PGRMC2 RNA and protein level in adrenals and ovaries of female *Aaas* KO mice.

Our group reported that female mice homozygous deficient for *Aaas* are infertile (Huebner et al., 2006). Carvalhal et al. recently presented that ALADIN is involved in mitotic and meiotic spindle assembly, chromosome segregation and production of fertile mouse oocytes (Carvalhal et al., 2015; Carvalhal et al., 2016). Interestingly, both PGRMC1 and 2 were shown to be involved in regulation of ovarian follicle development and therefore imply a neuroendocrine function (Wendler and Wehling, 2013). Deficiency of either MAPRs decreases the anti-apoptotic and anti-mitotic action of progesterone, although PGRMC2 seems to be important for anti-mitotic actions of the steroid. Depletion of either PGRMC1 or PGRMC2 leads to increased entry into cell cycle. Both proteins localise to the mitotic spindle and seem to exploit a distinct role during metaphase of mitosis, thereby suppressing entry into cell cycle. This effect is thought to be synergistic and does not seem to be additive (Griffin et al., 2014; Peluso et al., 2014; Sueldo et al., 2015).

Most recently, ALADIN and PGRMC2 have been identified to interact with the human centrosome-cilium interface (Gupta et al., 2015; Hanson et al., 2014; Yan et al., 2014). The centrosome is a fundamental organelle which participates in cell cycle progression and mitotic spindle assembly. Conclusively, ALADIN and PGRMC2 both seem to have an important role at the mitotic and meiotic spindle and to be involved in the sterility of female *Aaas* KO mice.

In summary, our straightforward work about the identification of the novel interactor of ALADIN, PGRMC2, provide new insights into the molecular function of the nucleoporin ALADIN in the pathogenesis of triple A syndrome. In the future it needs to be investigated how and why ALADIN associates with the microsomal protein at the perinuclear ER. In addition, their possible simultaneous role at the spindle apparatus shall await more research and reveal additional functions of ALADIN during cell division and meiosis.

## Materials and Methods

### Cell culture

NCI-H295R cells stably expressing GFP-ALADIN fusion protein or GFP were generated as described previously using the gamma-retroviral transfer vectors pcz-CFG5.1-GFP-*AAAS* and pcz-CFG5.1-GFP (Kind et al., 2009).

NCI-H295R cells transiently expressing PGRMC2-GFP fusion protein were generated as follows. Cells were transfected with pCMV6-AC-PGRMC2-GFP vector (RG204682) (OriGene Technologies, Rockville MD, USA) using X-tremeGENE HP DNA transfection reagent (Roche Diagnostics, Mannheim, Germany) following the manufacturer’s protocols. Cells were harvested or fixed after 48 hours.

In all exogenous expression models clones were selected by moderate expression of the desired fusion protein and true cellular localisation in order to exclude the possibility of false positive protein interactions.

Cells were cultured in DMEM/F12 medium (Lonza, Cologne, Germany) supplemented with 1 mM L-glutamine (Lonza, Cologne, Germany), 5% Nu-serum (BD Biosciences, Heidelberg, Germany), 1% insulin-tranferrin-selenium) (Gibco, Life Technologies, Darmstadt, Germany) and 1% antibiotic-antimycotic solution (PAA, GE Healthcare GmbH, Little Chalfont, United Kingdom).

NCI-H295R1-TR cells with *AAAS* knock-down or scrambled shRNA were generated, selected and cultured as described previously (Jühlen et al., 2015).

### Animals

All procedures were approved by the Regional Board for Veterinarian Affairs (AZ 24-9168.21-1-2002-1) in accordance with the institutional guidelines for the care and use of laboratory animals. C57BL/6J mice were obtained from Janvier Labs (Le Genest-Saint-Isle, France). *Aaas* KO mice were generated as described previously (Huebner et al., 2006).

### RNA extraction, cDNA synthesis and quantitative real-time PCR using TaqMan

Total RNA from cultured cells (n=10) and from frozen murine organs (at least four animals per genotype and sex) was isolated using the NucleoSpin RNA (Macherey-Nagel, Düren, Germany) according to the protocol from the manufacturer. Purity of the RNA was assessed using Nanodrop Spectrophotometer (ND-1000) (NanoDrop Technologies, Wilmington DE, USA). The amount of 500 ng of total RNA was reverse transcribed using the GoScript Reverse Transcription System (Promega, Mannheim, Germany) following the protocols from the manufacturer. Primers for the amplification of the target sequence were designed using Primer Express 3.0 (Applied Biosystems) and compared to the human or murine genome database for unique binding using BLAST search (National Center for Biotechnology Information, U.S. National Library of Medicine, 2013). The primer sequences are listed in the supplementary data of this article (**Table S3**).

The qPCR amplifications were performed in triplicates using the GoTaq Probe qPCR Master Mix (Promega) according to the manufacturer’s reaction parameter on an ABI 7300 Fast Real-Time PCR System (Applied Biosystems, Life Technologies, Darmstadt, Germany). In all results repeatability was assessed by standard deviation of triplicate C_t_s and reproducibility was verified by normalizing all real-time RT-PCR experiments by the C_t_ of each positive control per run.

### Immunoblots

After SDS-PAGE separation onto 4-12% PAGE (150 V for 1.5 hours) and electroblotting (30 V for 1.5 hours) (Invitrogen, Life Technologies, Darmstadt, Germany) onto Amersham hybond-ECL nitrocellulose membrane (0.45 μm) (GE Healthcare GmbH, Little Chalfont, United Kingdom) nonspecific binding of proteins to the membrane was blocked by incubation in PBS containing 3% BSA (Sigma-Aldrich, Munich, Germany) at room-temperature.

The membrane was then probed with primary antibodies either anti-ALADIN (B-11: sc-374073) (Santa Cruz Biotechnology, Inc., Heidelberg, Germany) (1:100 in 3% PBS/BSA) or anti-PGRMC2 (HPA041172) (Sigma-Aldrich, Munich, Germany) (1:200 in 5% PBS/milk powder) overnight at 4°C. Secondary antibodies goat anti-mouse IgG conjugated to horseradish peroxidase (1:2000 in 3% PBS/BSA) (Invitrogen, Life Technologies, Darmstadt, Germany) or goat anti-rabbit IgG conjugated to horseradish peroxidase (1:3000 in 5% PBS/milk powder) (Cell Signalling Technology Europe B.V., Leiden, Netherlands) were incubated one hour at room-temperature.

### Co-immunoprecipitation

For GFP co-IP lysates from NCI-H295R expressing GFP-ALADIN or PGRMC2-GFP were used. Lysates from cells expressing GFP were used as negative control. Cell lysates (500 μg protein) were added to the pre-equilibrated GFP-Trap_A agarose beads (ChromoTek GmbH, Planegg-Martinsried, Germany), gently resuspended by flipping the tube and bound over-night at constant mixing at 4°C. After washing steps the beads were gently re-suspended in 60 μl NUPAGE 2X LDS sample buffer and in order to dissociate the captured immunocomplexes from the beads, boiled at 95 °C for 10 minutes and Western Blot analysis was conducted with 20 μl of the eluate. The left 40 μl of the eluate using the lysates of NCI-H295R expressing GFP-ALADIN or GFP was further processed for proteomic profiling using mass spectrometry. These experiments following mass spectrometry analysis were repeated three times.

For co-IP of ALADIN or PGRMC2 lysates from NCI-H295R cells and Protein G UltraLink resin sepharose beads (Pierce, Thermo Scientific, Fischer Scientific, Schwerte, Germany) were used. Beads were gently resuspended in anti-ALADIN (2 μg/ml) or anti-PGRMC2 (HPA041172) (2 μg/ml) and as negative controls normal mouse or rabbit IgG (Invitrogen, Life Technologies, Darmstadt, Germany) (2 μg/ml). All antibodies were bound to the beads over-night at 4°C in a rotation chamber. After washing cell lysates (500 μg protein) were added to the beads, gently resuspended by flipping the tube and bound over-night as described before. After washing the beads were gently resuspended in 60 μl sample buffer containing dilution buffer, NUPAGE 1X LDS Sample Buffer and 1X Reducing Agent. The captured immunocomplexes were dissociated and the eluates were collected and processed by Western Blot as described previously. These experiments were repeated three times. The left 40 μl of the eluate after ALADIN co-IP and negative control was further processed for proteomic profiling using mass spectrometry. Mass spectrometry analysis was conducted once.

### Proteomic profiling using tandem mass spectrometry

Entire gel lanes were cut into 40 slabs, each of which was in-gel digested with trypsin (Shevchenko et al., 2006). Gel analyses were performed at the Mass Spectrometry Facility at the Max Planck Institute for Molecular Cell Biology and Genetics Dresden on a nano high-performance liquid chromatograph Ultimate interfaced on-line to a LTQ Orbitrap Velos hybrid tandem mass spectrometer as described previously (Vasilj et al., 2012).

Database search was performed against IPI human database (downloaded in July 2010) and NCBI protein collection without species restriction (updated in June 2014) using MASCOT software v.2.2. Scaffold software v.4.3.2 was used to validate MS/MS-based protein identifications. Protein probabilities were assigned by the Protein Prophet algorithm (Nesvizhskii et al., 2003).

### Immunofluorescence microscopy

Cells grown onto glass cover slips were fixed for 5 minutes with 4% PFA (SAV LP, Flinsbach, Germany) in PBS, permeabilised for 5 minutes with 0.5% Triton-X-100 in PBS and fixed again for 5 minutes. Blocking was performed for 30 minutes with 2% BSA/0.1% Triton-X-100 in PBS at room-temperature.

All antibodies used for immunofluorescence were diluted in blocking solution. Primary antibodies anti-ALADIN (1:25), or anti-PGRMC2 (HPA041172) (1:50) or anti-PGRMC2 (F-3: sc-374624) (Santa Cruz Biotechnology, Inc.) (1:25) and anti-NPC proteins (mAb414) (Covance, Berkley CA, USA) (1:800) were incubated at 4°C over-night in a humidified chamber. Secondary antibodies goat anti-mouse IgG Cy3 (1:800) (Amersham Biosciences, Freiburg, Germany), Alexa Fluor 488 and 555 goat anti-rabbit IgG (1:500) (Molecular Probes, Life Technologies) were incubated one hour at room-temperature in the dark.

Fluorescence was visualised using the confocal laser microscope TCS SP2 (Leica Microsystems, Mannheim, Germany). The experiments were repeated at least three times.

### Statistics of TaqMan analyses

Statistical analyses were made using the open-source software R version 3.3.0 and R Studio version 0.99.902 (R Core Team, 2015). Unpaired Wilcoxon-Mann-Whitney U-test was performed. During evaluation of the results a confidence interval alpha of 95% and P values lower than 0.05 were considered as statistically significant. Results are shown as box plots which give a fast and efficient overview about median, first and third quartile (25^th^ and 75^th^ percentile, respectively), interquartile range (IQR), minimal and maximal values and outliers.

## Acknowledgements

We thank Waldemar Kanczkowski for providing the NCI-H295R cells. Barbara Kind generously generated pseudo retroviruses containing pcz-CFG5.1-GFP-ALADIN and pcz-CFG5.1-GFP. We thank the Mass Spectrometry Facility at the Max Planck Institute for Molecular Cell Biology and Genetics Dresden for MS-based peptide analyses.

## Author contributions

RJ, AH and KK conceived and designed the experiments. RJ performed all experiments. DL helped with immunofluorescence staining and KK assisted with confocal microscopy. RJ analysed the data and wrote the paper. AH and KK helped improving the manuscript. All authors read the final version of the manuscript and gave their permission for publication.

